# Evaluating Large Language Models for Gene-to-Phenotype Mapping: The Critical Role of Full-Text Database Access

**DOI:** 10.1101/2025.06.11.659165

**Authors:** Nicolas Matthew Suhardi, Anastasia Oktarina, Julia Retzky, Damanpreet Dhillon, Dona Ninan, Mathias P.G. Bostrom, Xu Yang, Vincentius Jeremy Suhardi

**Author notes:** Corresponding author, Email address (Vincentius Jeremy Suhardi).

## Abstract

Transformer-based large language models (LLMs) have demonstrated significant potential in the biological and medical fields due to their ability to effectively learn from large-scale, diverse datasets and perform a wide range of downstream tasks. However, LLMs are limited by issues such as information processing inaccuracies and data confabulation. These limitations hinder their utility for literature searches and other tasks requiring accurate and comprehensive extraction of information from extensive scientific literature. In this study, we evaluated the performance of various LLMs in accurately retrieving peer-reviewed literature and mapping correlations between 198 genes and six phenotypes: bone formation, cartilage formation, fibrosis, cell proliferation, tendon formation, and ligament formation. Our analysis included three types of models. First, standard transformer-based LLMs (ChatGPT-4o and Gemini 1.5 Pro). Second, specialized LLMs with dedicated custom databases containing peer-reviewed articles (SciSpace and ScholarAI). Third, specialized LLMs without dedicated databases (PubMedGPT and ScholarGPT). Using human-curated gene-to-phenotype mappings as the ground truth, we found that specialized LLMs with dedicated databases achieved the highest accuracy (>80%) in gene-to-phenotype mapping. Additionally, these models were able to provide relevant peer-reviewed publications supporting each gene-to-phenotype correlation. These findings underscore the importance of database augmentation and specialization in enhancing the reliability and utility of LLMs for biomedical research applications.

## Introduction

Understanding the relationship between genes and their cellular phenotypes is crucial in the biomedical field. It provides insights into how genetic perturbations impact cellular behavior[1]. This knowledge is essential for determining how gene function relates to cellular phenotypes. The process of mapping genetic perturbations to phenotypes typically involves two approaches. The first approach identifies the gene(s) associated with a specific observed phenotype. The second approach evaluates the phenotypic changes resulting from specific gene perturbations. Accurate correlations between genes and phenotypes are foundational for various applications, including cellular type classification in single-cell omics analysis[2]. However, the traditional approach of perturbing one gene at a time is labor-intensive and time-consuming. The vast number of genes within the genome, coupled with the virtually limitless combinations of phenotypes resulting from perturbation of a single gene—and further compounded by the exponential growth of research publications—has made the task of accurately mapping genes to phenotypes increasingly challenging[3].

Given these limitations of traditional approaches, large language models (LLMs), particularly transformer-based architectures, have garnered significant interest in biology and medicine. Transformer-based LLMs effectively learn from extensive and diverse datasets, facilitating a wide range of downstream applications. These models frequently surpass the performance of task-specific models trained from scratch [4, 5]. Specifically, LLMs have demonstrated substantial utility in elucidating complex relationships within genomic (DNA sequences), transcriptomic (RNA sequences), and proteomic (protein sequences and structures) datasets [6]. Central to transformer architectures is the self-attention mechanism [7], which enables the model to capture context and manage long-range dependencies more effectively than recurrent neural networks (RNNs) or long short-term memory (LSTM) models [8]. Moreover, the multi-head attention mechanism inherent in transformers allows parallel processing of information, significantly enhancing computational efficiency and reducing processing time relative to RNNs and LSTMs[7].

Despite their significant potential and growing use in biomedical research, large language models (LLMs) have notable limitations. Previous studies have underscored issues concerning their accuracy [9] and tendency to generate false or misleading information, often termed hallucinations [10]. Additionally, the lack of transparency regarding training data and inherent biases can perpetuate errors and misinformation [11]. Given the expanding use of LLMs in biomedical research, particularly for mapping gene-phenotype relationships[2, 12], it is essential to assess their accuracy rigorously in this specific task.

To address these concerns and evaluate LLM performance in the critical area of gene-phenotype identification, we assessed transformer-based LLMs—specifically ChatGPT-4o and Gemini 1.5 Pro. These models were evaluated based on their embedded biological knowledge, as well as their ability to leverage information from PubMed and other open-access literature sources. We utilized six key phenotypes as representative proxies: cell proliferation, bone formation, fibrosis, cartilage formation, tendon formation, and ligament formation. Human-curated, peer-reviewed literature searches served as our benchmark for accuracy.

Both ChatGPT-4o and Gemini 1.5 Pro demonstrated moderate accuracy in identifying gene-phenotype relationships. Importantly, granting specialized access to abstracts and full-length articles significantly improved the accuracy of these models. These findings suggest that transformer-based LLMs, when specialized with direct literature access, hold promise as reliable tools for accurately mapping gene-phenotype relationships.

## Methods

### Transformer-Based Large Language Models

Publicly available transformer-based large language models (LLMs) ChatGPT-4o (“o” for “omni”) and Gemini 1.5 Pro were used through their user interface. As of May 2025, ChatGPT-4o had around 1.76 trillion parameters [13] and has a context window of 128,000 tokens. Gemini 1.5 Pro has a context window of 2 million tokens, though the exact number of model parameters has not been officially disclosed by Google. Both ChatGPT-4o and Gemini 1.5 Pro were able to access metadata of PubMed abstracts but unable to directly access the PubMed database.

### Model Access and Implementation

All models evaluated in this study are closed-source models and were accessed through their respective web interfaces without any fine-tuning, training, or modification. Each model was used in its standard, publicly available configuration through browser-based interfaces. No custom code was developed for this study; all data collection and analysis were performed manually using spreadsheet software.

### Data Collection Methodology

This study employed a completely manual approach to data collection and analysis to ensure reproducibility and transparency. The entire experimental protocol was conducted without automated scripts or custom software development. The following step-by-step methodology was used:

Query Execution: For each of the 1,188 gene-phenotype combinations per model, queries were systematically submitted through the web interfaces of the respective large language models (LLMs). Each query adhered rigorously to the standardized prompt format detailed in the LLM Prompts section. Queries were individually executed via manual entry into browser-based interfaces, ensuring consistent methodology for each gene-phenotype combination.

Response Collection: Responses generated by the LLMs were systematically collected by manually transferring data from the web interface to Microsoft Excel spreadsheets. Separate spreadsheets were maintained for each LLM model evaluated. Each spreadsheet entry comprehensively recorded the gene name, associated phenotype, the full LLM response, categorized outcomes (“increase,” “decrease,” or “inconclusive”), provided citations, and relevant journal impact factor information.

Reference Verification: URLs provided by the LLMs linking to peer-reviewed publications were individually verified through direct manual access. Each URL was critically evaluated to ensure it directed accurately to the referenced publication, confirmed congruence with the publication title and authorship details provided by the LLM, and assessed its relevance to the specific gene-phenotype relationship queried. URLs or reference titles that led to non-existent publications were categorized as “hallucination.” URLs or reference titles that correctly directed to a publication but were unrelated to the queried gene-phenotype relationship were categorized as “irrelevant.”

### Scientific-Focused Large Language Models

For scientific-focused large language models (LLMs), three types of GPT-based systems were utilized:

1. LLMs with access to dedicated databases of full-length articles: These include models like GPT-4-based SciSpace (Typeset.io) and ScholarAI.
2. LLMs without dedicated databases but with direct access to publicly available sources: Examples include
3. GPT-4-based ScholarGPT, which can access various scholarly databases and academic search engines.
4. LLMs without access to full-length articles, customized to draw information from abstracts: An example is PubMedGPT, which primarily utilizes abstracts from the PubMed database.

#### Technical Implementation

These specialized models are implemented as closed-source, publicly available, custom GPTs. Each model utilizes the ChatGPT platform’s internet search capabilities and custom API integrations as tools. SciSpace and ScholarAI employ API calls to their respective proprietary databases and search systems, while ScholarGPT uses API connections to established scholarly indexes and journal databases. PubMedGPT operates without external API calls, relying on its specialized instructions to extract information from its training data focused on PubMed abstracts. The base models (ChatGPT-4o and Gemini 1.5 Pro) utilize the standard internet search tools provided by their respective platforms.

#### GPT-4-based SciSpace

SciSpace’s general knowledge cutoff was October 2023. However, it can retrieve, analyze, and synthesize information from its proprietary database of over 300 million research papers. Through its integration with this database, SciSpace provides real-time access to research papers up to 2025. Additionally, SciSpace can access publicly available databases, including CrossRef, PubMed, arXiv, Springer, Elsevier, IEEE Xplore, Google Scholar, and patent databases.

#### GPT-4-based ScholarAI

Like SciSpace, ScholarAI’s general knowledge cutoff was October 2023. It has access to the dedicated ScholarAI database, which contains more than 200 million research papers, and can also draw from publicly available sources such as CrossRef, PubMed, arXiv, Springer, Elsevier, IEEE Xplore, Google Scholar, and patent databases.

#### GPT-4-based ScholarGPT

ScholarGPT had a general knowledge cutoff in October 2023 and lacked a dedicated database. Instead, it relies on real-time access to abstracts and open-access full-text articles from various publicly available scholarly databases and academic search engines.

#### PubMedGPT

PubMedGPT was not trained on full-text manuscripts from PubMed or other databases. Its training included publicly available data up to 2023. PubMedGPT is designed to provide citations from PubMed and integrate findings from multiple studies to synthesize conclusions.

### Queries

Six phenotypes were utilized in this study: bone formation, fibrosis, cartilage formation, cell proliferation, tendon formation, and ligament formation. These phenotypes were chosen as representative topics of interest for skeletal and connective tissue researchers, reflecting both mineralized and soft tissue outcomes relevant to musculoskeletal biology.

A power analysis was conducted to determine the minimum number of genes required to detect at least a 20% difference with a power of 80% and a significance level (p-value) of 0.05. The analysis indicated that at least 91 genes would be needed. To ensure robust statistical power and comprehensive coverage of biological pathways, we expanded our dataset to 198 genes.

To ensure comprehensive representation, 198 genes previously reported in peer-reviewed studies to impact at least one of the six phenotypes were selected as genes of interest (Table 1). These 198 genes were selected to encompass various genetic pathways. These include bone morphogenetic pathway (BMP Pathway), wingless/integrated pathway (WNT Pathway), transforming-growth factor beta pathway (TGFβ pathway), Hedgehog pathway, NF-kB pathway, fibroblast growth factor pathway (FGF pathway), and additional pathways relevant to connective tissue biology. The expanded gene set includes established markers of skeletal development (e.g., RUNX2, SOX9, COL1A1), tissue remodeling factors (e.g., MMP13, ADAMTS5), mechanosensitive channels (e.g., PIEZO1, PIEZO2), and tissue-specific transcription factors (e.g., SCX for tendon, GDF5 for cartilage). For each gene, six distinct LLM queries were created, corresponding to the six phenotypes of interest: bone formation, fibrosis, cartilage formation, cell proliferation, tendon formation, and ligament formation. This resulted in 1,188 total queries per LLM model tested.

**Table 1:** List of genes used in this study.

| **Phenotype** | **Gene Names** |
| --- | --- |
| **Fibrosis** | Piezo1, En1, Bglap, Osterix, Wnt1, Wnt3a, Osteopontin, Pth1R, LRP5, LRP6, RSPO1, RSPO2, RSPO3, Sclerostin, Dkk1, Dkk2, Piezo2, Spry2, Dkk3, Dkk4, Gli1, Gli2, Gli3, BMP1, BMP2, Col2a1, Prx1, Sox2, WIF-1, Sp7, Embigin |
| **Bone** | Alp, Col1a1, Runx2, Krm1, Smad2, Smad4, Smad3, Runx1, IGF-1, IGF1R, Slit3, Shn3, Ostelectin, Fgf2, Fgfr2, Fgf23, Acta2, Ikbkb, Lats2, Galectin1, Pdgfra, Pdgfrb, Tgfr2, Asxl1, Cxcl4, Cxcr2, Smpd3, Sox9, Nfkb1, Ctgf, Timp1, Pkd1, Tnfrsf11b, Tnfa, Bmp7 |
| **Cartilage** | Plcg1, Alk1, Alk5, Smad1, Smad8, Pinch1, Pinch2, Mek1, Mek2, Tgfr1, CCn2, Fgfr1, Mef2c, Epas1, Msx2, Sox5, Sox6, Ddr2, Scn9a, Erg, Rela, Mmp13, Adamts5, Prg4, Gdf5, Frzb, Ctnnb, Egfr, Nfatc2, Nfatc1, Egr1, Sparc, Sost |
| **Cell Proliferation** | Grem1, Trpv4, Creb5, Notch2, MKX, EGR2, HIF1A, NRF2, SIRT1, TSC1, AKT1, CAV1, DICER1, ESR1, GDF6, GDF7, TGF-β2, TGF-β3, Noggin, ACVR1, SMO, IL-4, IL-10, IL-13, IL-17A, IL-33, CCL2, CXCL12, CXCR4, TLR4, NOS2, NOS3 |
| **Tendon Formation** | SCX, HO-1, FLII, SFRP2, Substance P, COX-2, LGALS3, PPARG, COL3A1, COL5A1, COL10A1, COL12A1, COL14A1, DCN, BGN, FMOD, LUM, VCAN, TNC, POSTN, THBS1, THBS2, THBS3, THBS4, COMP, MGP, FN1, CLEC3B, TSG-6, IGFBP5, CDH11, MMP3, MMP8, MMP9 |
| **Ligament Formation** | MMP10, MMP12, MMP14, MMP2, ADAMTS2, LOX, LOXL2, PLG, PLAU, SERPINE1, MKL1, YAP1, WWTR1, SMAD7, ILK, ITGB1, ITGA5, PDGFB, VEGFA, ANGPT1, ANGPT2, CDKN1A, CDKN2A, TGFB1I1, PTEN, MAPK14, MAPK1, MAPK3, MAPK8, RELB, STAT3, IGF2, TIMP2 |

### LLM Prompts

One-shot prompting approach was utilized to ensure consistent and structured responses across all models. The prompt design included model-specific instructions to direct each LLM to use its appropriate database or search capabilities, followed by a standardized query structure and output format.

The prompt consisted of three main components. First, a model-specific instruction directing the LLM to use its designated database or search tool. Second, the core query asking whether a specific gene increases, decreases, or has an inconclusive relationship with a given phenotype based on peer-reviewed literature. Third, detailed formatting instructions requiring a single-word answer followed by supporting references and impact factor information.

To enhance response consistency and accuracy, an example of the expected output format was included as port of the prompt (**Figure 1**). The one-shot example was included to allow the LLM to return consistent structured answer: a bracketed single-word answer categorized as increase, decrease, or inconclusive, followed by numbered reference links, and the highest impact factor among cited journals.

**Figure 1.**
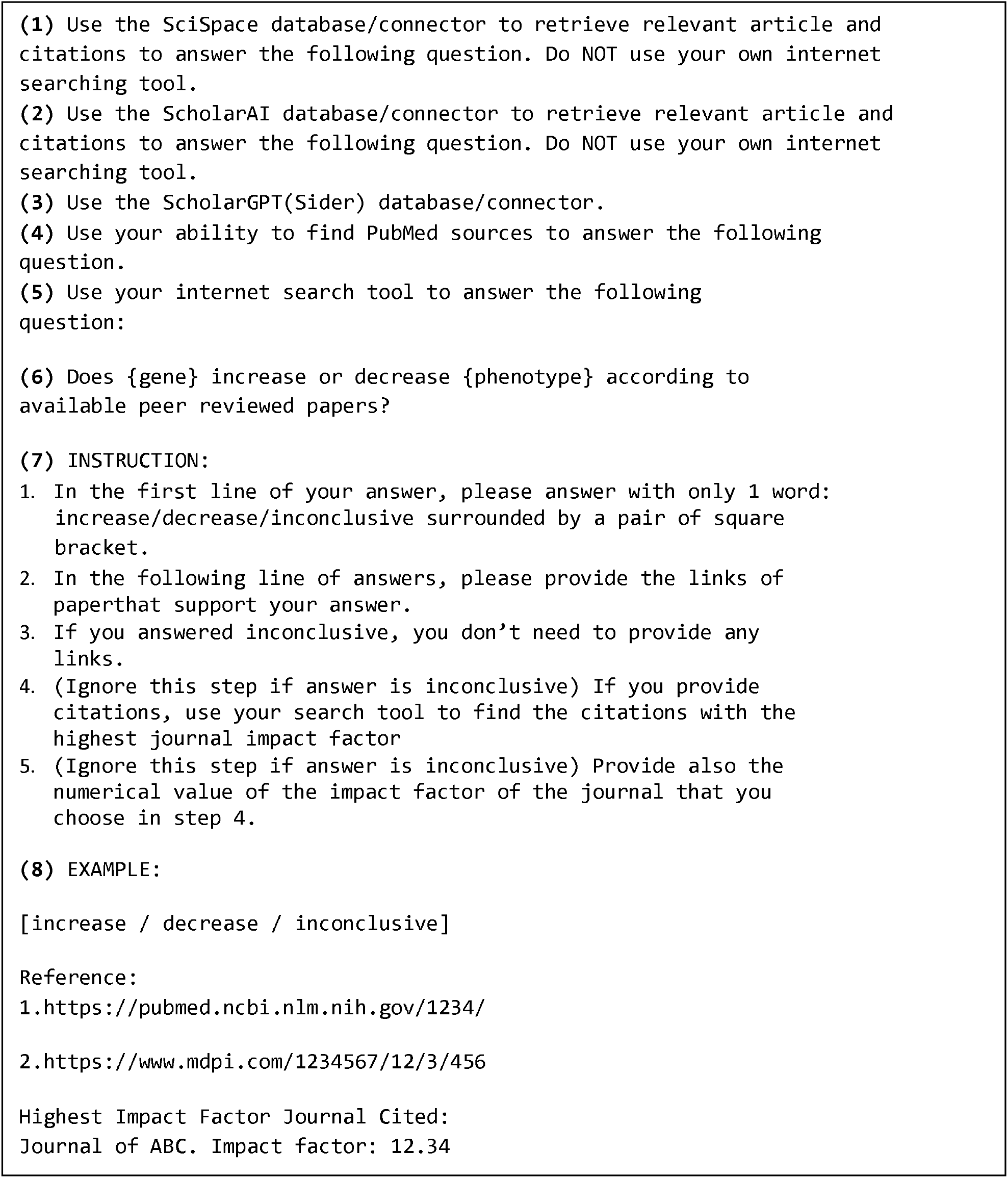
Standardized prompt template used for querying large language models. The prompt consists of three main components: (1) Model-specific instructions (part 1-5) directing each LLM to use its designated database or search capabilities, with only the relevant instruction provided to each model; (2) Core query structure (part 6) following a standardized format asking whether a specific gene increases, decreases, or has an inconclusive relationship with a given phenotype based on peer-reviewed literature; and (3) Detailed response formatting instructions (part 7) requiring a single-word answer in square brackets followed by supporting references and impact factor information. The example output (part 8) demonstrates the expected response format to ensure consistency across all 1,188 queries per model. This structured one-shot prompting approach minimized response variability and facilitated systematic data extraction across all six LLM models tested.

All links to peer-reviewed manuscripts provided by the LLM prompts were manually verified to ensure they correspond to the intended and relevant manuscripts.

### Reference Quality Assessment

To rigorously evaluate the quality and reliability of references provided by each LLM, we employed four specific metrics:

1. **Accuracy Assessment:** Responses for each gene-phenotype query were assessed against ground truth determinations established by manual curation of peer-reviewed literature by the senior author (VJS). A response was considered accurate if the LLM’s determination (increase, decrease, or inconclusive) matched the manually curated classification.
2. **Topic-Irrelevant Citations:** For queries in which references were provided, we assessed relevance to both the specified gene and the phenotype queried. A citation was categorized as topic-irrelevant if the cited paper did not contain information directly supporting the gene-phenotype relationship. The topic-irrelevant citation rate was calculated as the proportion of queries where all provided references failed to substantiate the claimed gene-phenotype association.
3. **Hallucination:** References provided by LLMs were verified to determine if they corresponded to actual publications. A reference was classified as hallucination if: (a) the provided URL directed to a non-existent page or a different publication than indicated, or (b) the title and authors listed by the LLM did not match the actual publication accessible through the URL. The hallucination rate was calculated as the proportion of queries where all cited references were either non-existent or inaccurately represented.
4. **Impact Factor Analysis:** The journal impact factor was recorded for all valid peer-reviewed references using the most recent Journal Citation Reports®. For queries providing multiple valid references, we utilized the highest impact factor among cited journals. Queries resulting in inconclusive determinations or lacking valid peer-reviewed citations were assigned an impact factor of zero.

### Statistical Analysis

Logistic regression was performed to analyze the performance of large language models (LLMs) in providing correct answers, using human responses as the ground truth. ChatGPT-4o (base model) was treated as the reference model. The p-value was calculated to test the null hypothesis that the log-odds difference between the tested model and the reference model is zero (i.e., the performance of the tested model is the same as that of the reference model). Statistical analysis was conducted using GraphPad Prism v9.5, with a significance threshold set at p<0.05.

## Results

### ChatGPT-4o and Google Gemini 1.5 Pro demonstrated comparable moderate accuracy in mapping genes to phenotype

We investigated the performance of an OpenAI-based large language model (LLM) (ChatGPT-4o) in comparison to the Gemini 1.5 Pro LLM to map genes to phenotype based on peer-reviewed literature. To evaluate the ability of the LLMs to map genes to phenotypes, each model was tasked with mapping 198 genes (as described in the Methods section) to six phenotypes: bone formation, cartilage formation, fibrosis, cell proliferation, tendon formation, and ligament formation. This resulted in a total of six queries per gene and 1,188 queries per LLM. Manual mapping of genes to phenotypes by the senior author (VJS), based on peer-reviewed journal articles, was used as the ground truth for comparison.

ChatGPT-4o demonstrated significantly higher accuracy in gene-to-phenotype mapping compared to Gemini 1.5 Pro (59.3% vs. 55.8%, respectively; p = 0.04, **Figure 2a**). Although prompts explicitly requested references from peer-reviewed articles, not all queries yielded relevant citations. Notably, ChatGPT-4o provided a significantly greater proportion of irrelevant citations compared to Gemini 1.5 Pro (18.4% vs. 5.5%, respectively; p = 0.0001, **Figure 2b**). Furthermore, ChatGPT-4o exhibited a higher hallucination rate than Gemini 1.5 Pro (16.4% vs. 7.1%, respectively; p = 0.0001, **Figure 2b**). Performance varied across the six queried phenotypes, with both models achieving their highest accuracy in fibrosis-related queries (ChatGPT-4o: 73.2%, Gemini 1.5 Pro: 69.2%, **Supplementary Figure 1**). Conversely, the lowest accuracy was observed in cartilage formation queries for ChatGPT-4o (45.9%) and ligament formation queries for Gemini 1.5 Pro (44.4%).

**Figure 2.**
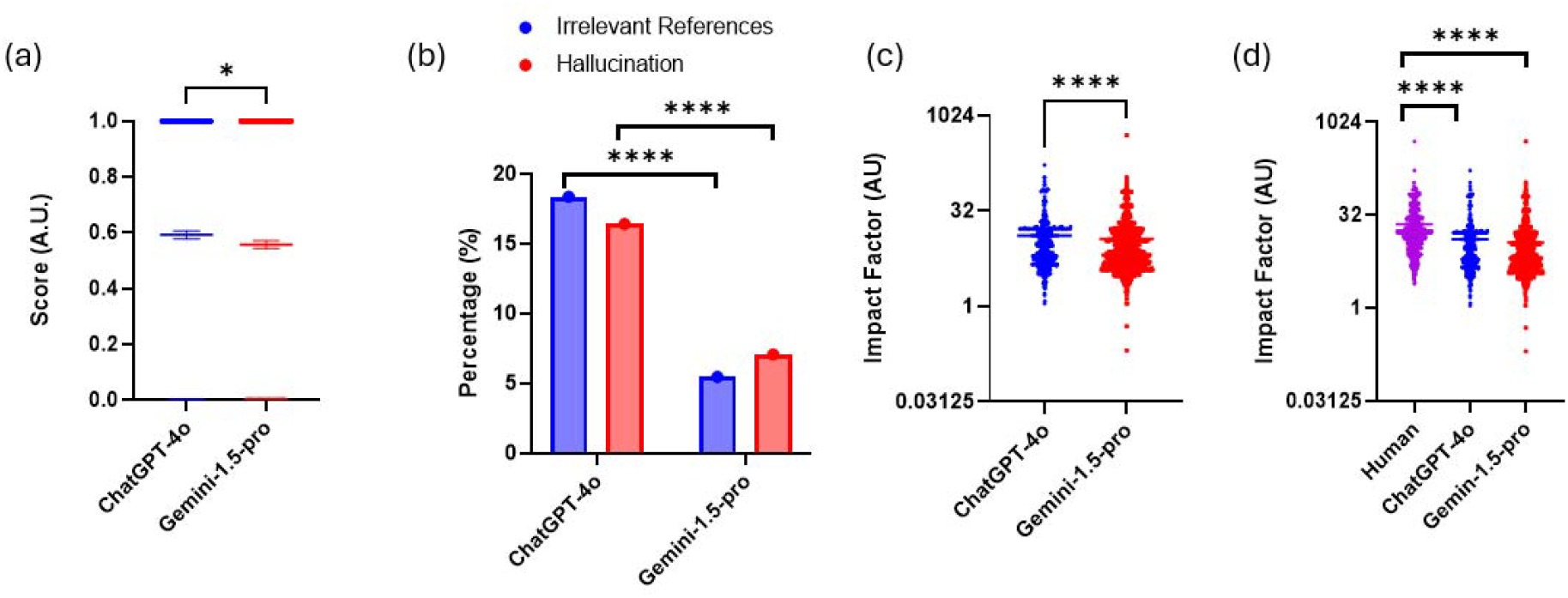
Accuracy and impact factor of references provided by ChatGPT4o and GeminiAI. (a) Accuracy of correct gene-to-phenotype mappings by ChatGPT4o and GeminiAI. Each dot represents a single gene-to-phenotype query, with a value of 1 indicating agreement between the tested LLM and human answers, and 0 indicating a discrepancy. Data are presented as mean ± s.e.m., analyzed using the Wilcoxon pair-matched signed-rank test. (b) Proportion of hallucinations (red bars) and irrelevant references (blue bars) provided by ChatGPT4o and GeminiAI. Data are presented as mean ± s.e.m., analyzed using the Chi-square test.(c-d) Impact factor of references provided by human, ChatGPT4o, and GeminiAI. Each dot represents the impact factor of a single gene-to-phenotype query. Data are presented as mean ± s.e.m., analyzed using the Wilcoxon pair-matched signed-rank test. *p<0.05, **p<0.01, ***p<0.001, ****p<0.0001.

Among relevant peer-reviewed articles, the Wilcoxon paired signed-rank test indicated significantly higher impact factors per correct answer between ChatGPT-4o and Gemini 1.5 Pro (14.6 vs. 14.0, respectively; p = 0.001, p=0.0001, **Figure 2c**). Human-provided references had a statistically significantly higher impact factor than both ChatGPT-4o and Gemini 1.5 Pro (p = 0.0001 for human vs. ChatGPT-4o, and p = 0.0001 for human vs. Gemini 1.5 Pro; **Figure 2d**).

### The scientific-focused GPT-based LLM without its own database underperformed the base GPT-based LLM in mapping genes to phenotype

Building on these baseline comparisons between standard LLMs, we investigated whether LLMs specialized for scientific purposes could improve gene-phenotype mapping performance. To address this question, we utilized publicly available ChatGPT-4o-based, scientifically specialized LLMs, including PubMedGPT, SciSpace, ScholarGPT, and ScholarAI. These scientific LLMs were categorized into three main groups. First, LLMs with access to dedicated databases of full-length articles (SciSpace and ScholarAI). Second, LLMs without dedicated databases but with direct access to publicly available sources (ScholarGPT). Third, LLMs without access to full-length articles, customized to extract information from PubMed abstracts (PubMedGPT).

PubmedGPT, an LLM customized to extract information from PubMed abstracts, demonstrated similar accuracy compared to ChatGPT-4o (57.6% for PubmedGPT vs 59.3% for ChatGPT-4o, p = 0.23; **Figure 3a**). PubmedGPT had a comparable hallucination rate compared to ChatGPT-4o (16.4 % vs 16.2%, respectively; **Figure 3b**). However, PubmedGPT had a significantly lower percentage of irrelevant references compared to ChatGPT-4o (5.4% vs. 18.3%, respectively; p = 0.0001; **Figure 3b**). Among the relevant peer-reviewed articles and using the Wilcoxon pair-matched signed rank test, the impact factor of references provided by PubmedGPT had significantly lower impact factors to those from ChatGPT-4o and human (11.6 ± 11.9 for PubmedGPT, 13.1 ± 15.3 for ChatGPT-4o, 22.9 ± 25.0 for human, p = 0.02 for PubmedGPT vs ChatGPT-4o and p=0.0001 for PubmedGPT vs Human; **Figure 3c-d**). The phenotype-specific analysis revealed that despite its specialization in biomedical abstracts, PubMedGPT showed consistently low accuracy across all phenotypes (42.4-65.2% range), with particularly poor accuracy in cartilage formation (42.4%, **Supplementary Figure 1**).

**Figure 3.**
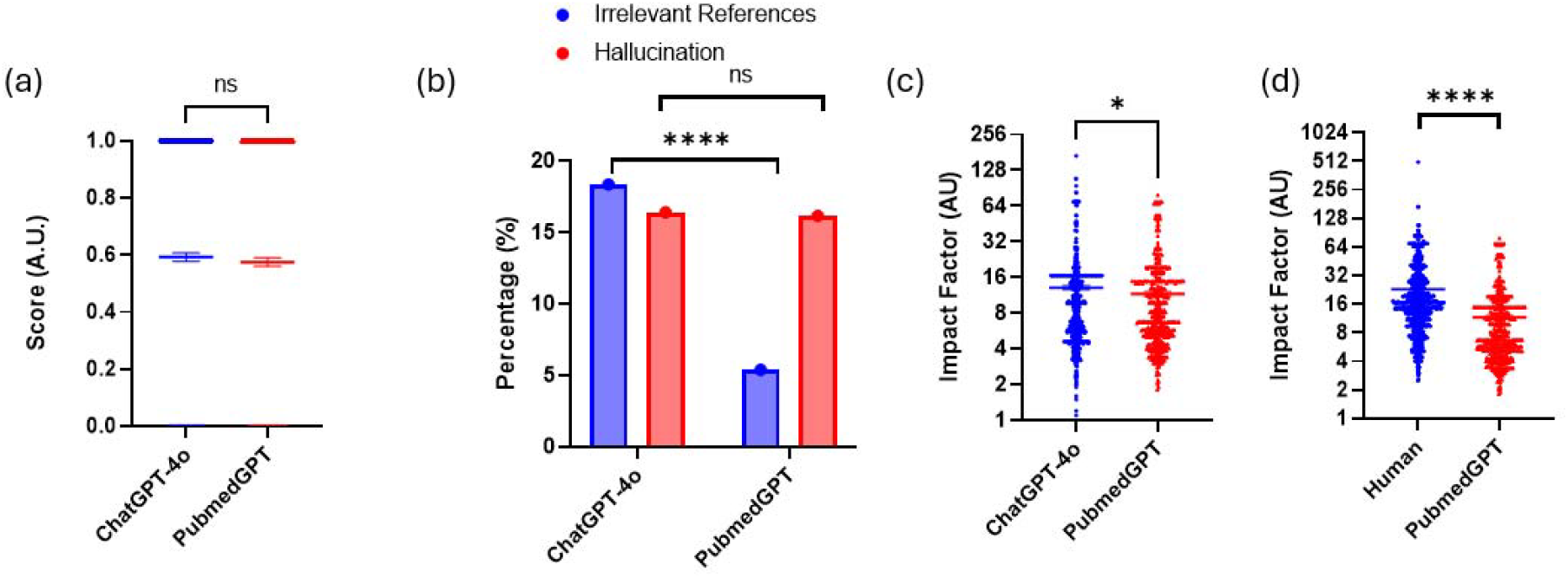
Accuracy and impact factor of references provided by PubmedGPT. (a) Accuracy of gene-to-phenotype mappings by ChatGPT4o and PubmedGPT. Each dot represents a single gene-to-phenotype query, with a value of 1 indicating agreement between the tested LLM and human answers, and 0 indicating a discrepancy. Data are presented as mean ± s.e.m., analyzed using the Wilcoxon pair-matched signed-rank test. (b) Proportion of hallucinations (red bars) and irrelevant references (blue bars) provided by ChatGPT4o and PubmedGPT. Data are presented as mean ± s.e.m., analyzed using the Chi-square test.(c-d) Impact factor of references provided by human, ChatGPT4o, and PubmedGPT. Each dot represents the impact factor of a single gene-to-phenotype query. Data are presented as mean ± s.e.m., analyzed using the Wilcoxon pair-matched signed-rank test. *p<0.05, **p<0.01, ***p<0.001, ****p<0.0001.

### ScholarGPT with public database access showed limited improvement over base ChatGPT-4o

To further explore the role of database access beyond abstract-only models, we investigated whether GPT-based LLMs with direct access to publicly available scholarly sources could improve performance. We compared the ability of ScholarGPT to the base ChatGPT-4o and human to map genes to phenotype based on relevant peer-reviewed articles.

ScholarGPT had comparable accuracy compared to ChatGPT-4o (58.6 % for ScholarGPT vs 59.3 % for ChatGPT-4o, p = 0.064; **Figure 4a**). Compared to ChatGPT-4o, ScholarGPT demonstrated significantly lower hallucination rate (5.9% for ScholarGPT vs 16.4% for ChatGPT-4o, p=0.0001, **Figure 4b**). ScholarGPT provided significantly lower irrelevant reference, with only 2.1% topic-irrelevant citations, significantly lower than ChatGPT-4o’s 18.4% (p = 0.0001). Among the relevant peer-reviewed articles provided and using the Wilcoxon pair-matched signed rank test, the impact factor of references provided by ScholarGPT were comparable to the base ChatGPT-4o (13.1 ± 15.3 for ChatGPT-4o, 13.4 ± 14.3 for ScholarGPT, p = 0.35, **Figure 4c**) but significantly lower compared to human (13.4 ± 14.3 ScholarGPT and 22.9 ± 25.0 for human, p= 0.0001, **Figure 4d**). The phenotype-specific heatmap demonstrated that ScholarGPT maintained consistent performance across all phenotypes with moderate accuracy (54.5-72.7%, **Supplementary Figure 1**).

**Figure 4.**
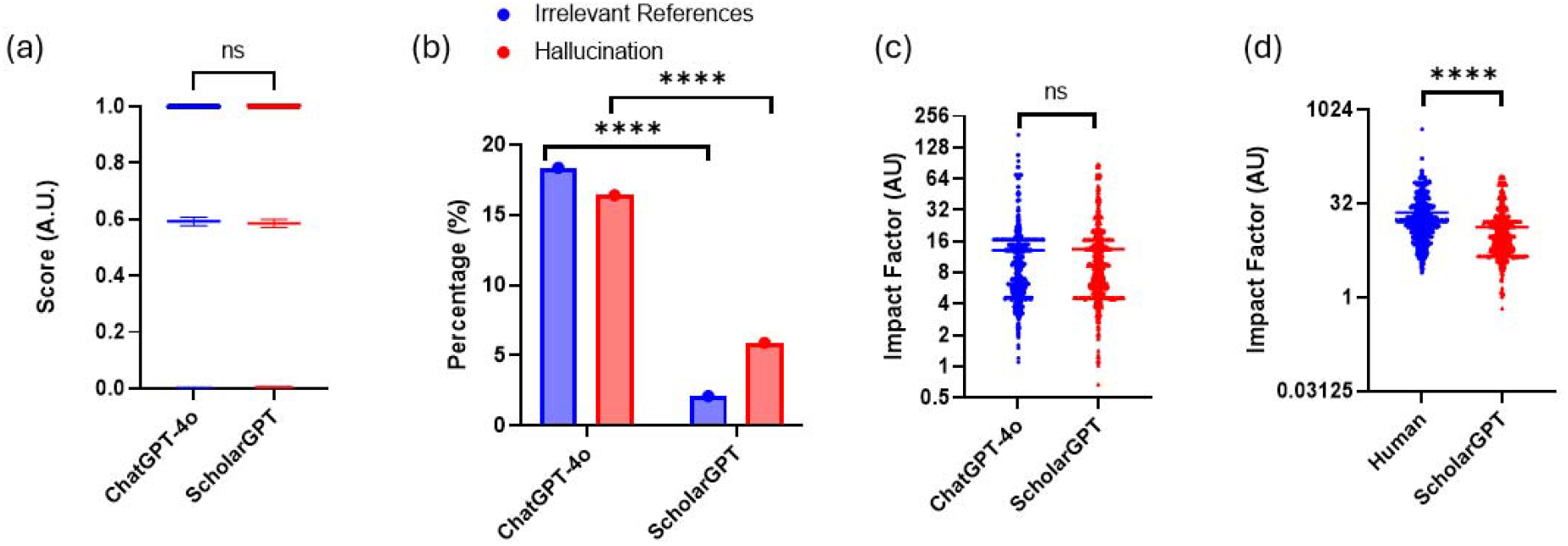
Accuracy and impact factor of references provided by ScholarGPT. (a) Accuracy of gene-to-phenotype mappings by ChatGPT4o and ScholarGPT. Each dot represents a single gene-to-phenotype query, with a value of 1 indicating agreement between the tested LLM and human answers, and 0 indicating a discrepancy. Data are presented as mean ± s.e.m., analyzed using the Wilcoxon pair-matched signed-rank test. (b) Proportion of hallucinations (red bars) and irrelevant references (blue bars) provided by ChatGPT4o and ScholarGPT. Data are presented as mean ± s.e.m., analyzed using the Chi-square test.(c-d) Impact factor of references provided by human, ChatGPT4o, and ScholarGPT. Each dot represents the impact factor of a single gene-to-phenotype query. Data are presented as mean ± s.e.m., analyzed using the Wilcoxon pair-matched signed-rank test. *p<0.05, **p<0.01, ***p<0.001, ****p<0.0001.

### The scientific-focused GPT-based LLMs with their own dedicated databases outperformed the base GPT LLM in mapping genes to phenotype

We next assessed whether large language models (LLMs) integrated with dedicated databases could enhance performance in gene-phenotype mapping tasks. Models augmented with dedicated databases containing full-length articles, specifically SciSpace and ScholarAI, exhibited significantly improved accuracy compared to ChatGPT-4o (74.9% for SciSpace vs. 59.3% for ChatGPT-4o, p = 0.0001; 81.0% for ScholarAI vs. 59.3% for ChatGPT-4o; p=0.0001, **Figure 5a**). Additionally, these database-enhanced models outperformed other LLMs lacking dedicated databases, including those relying solely on publicly available resources (e.g., ScholarGPT) and those tailored specifically for retrieving information from PubMed abstracts (e.g., PubMedGPT) (**Figure 5b**).

**Figure 5.**
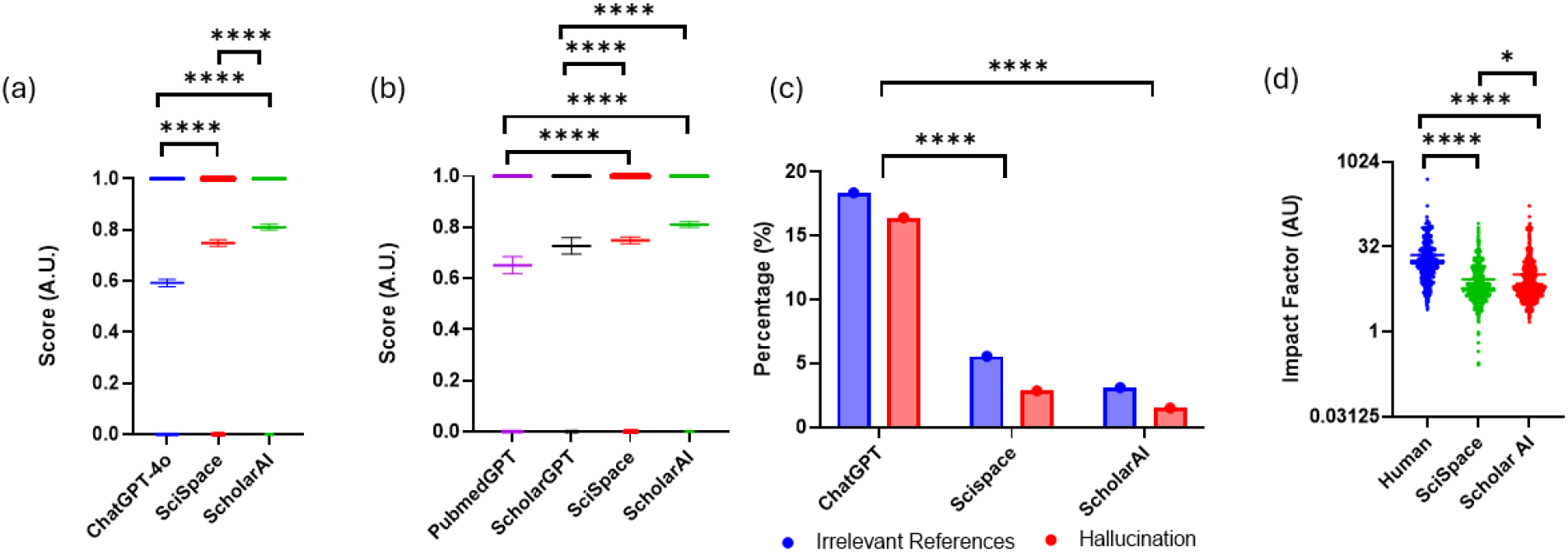
Accuracy and impact factor of references provided by SciSpace and ScholarAI. (a) Accuracy of gene-to-phenotype mappings by ChatGPT4o, SciSpace, and ScholarAI. Each dot represents a single gene-to-phenotype query, with a value of 1 indicating agreement between the tested LLM and human answers, and 0 indicating a discrepancy. Data are presented as mean ± s.e.m., analyzed using the Wilcoxon pair-matched signed-rank test. (b) Percentage of correct gene-to-phenotype mappings by PubmedGPT, ScholarGPT, SciSpace, and ScholarAI. Each dot represents a single gene-to-phenotype query, with a value of 1 indicating agreement between the tested LLM and human answers, and 0 indicating a discrepancy. Data are presented as mean ± s.e.m., analyzed using the Wilcoxon pair-matched signed-rank test. (c) Proportion of hallucination (red bars) and irrelevant references (blue bars) provided by ChatGPT4o, SciSpace, and ScholarAI. Data are presented as mean ± s.e.m., analyzed using the Chi-square test. (d) Impact factor of references provided by human, ChatGPT4o, SciSpace, and ScholarAI. Each dot represents the impact factor of a single gene-to-phenotype query. Data are presented as mean ± s.e.m., analyzed using the Wilcoxon pair-matched signed-rank test. *p<0.05, **p<0.01, ***p<0.001, ****p<0.0001.

Moreover, SciSpace and ScholarAI demonstrated significantly lower rates of hallucination compared to ChatGPT-4o (1.5% for ScholarAI, 2.86% for SciSpace, and 16.4% for ChatGPT-4o; p = 0.0001; **Figure 5c**). Similarly, both SciSpace and ScholarAI showed markedly lower rates of topic-irrelevant citations relative to ChatGPT-4o (3.1% for ScholarAI, 5.55% for SciSpace, and 18.3% for ChatGPT-4o; p = 0.0001; **Figure 5c**).

The average impact factors of references cited by ScholarAI were significantly higher than those cited by SciSpace, although both were notably lower compared to human-provided references (10.3 ± 12.7 for ScholarAI, 8.5 ± 8.0 for SciSpace, and 22.9 ± 25.0 for human; p = 0.0016 for ScholarAI vs. SciSpace; p = 0.001 for human vs. SciSpace and human vs. ScholarAI; **Figure 5d**). The phenotype-specific heatmap revealed that ScholarAI and SciSpace achieved remarkably consistent high performance across all phenotypes (79.3-82.3% for ScholarAI and 71.2-80.8% for SciSpace, **Supplementary Figure 1**).

## Discussion

Accurate gene-to-phenotype mapping at cellular and tissue levels is critical for single-cell omics analyses[14], elucidating disease correlations[15], assessing genetic perturbations[2], and advancing the mechanistic understanding of disease pathophysiology[16]. Currently, manual human curation from peer-reviewed literature remains the gold standard. However, given the exponential growth in scientific publications, there is a pressing need for automated or minimally supervised methods capable of efficiently handling this task.

An ideal automated system for gene-to-phenotype mapping must meet three critical requirements: (1) accurate retrieval of relevant literature, (2) precise extraction and synthesis of relevant information, and (3) generation of reliable gene-to-phenotype correlations from aggregated data. Meeting these requirements would markedly enhance efficiency and accuracy while significantly reducing reliance on manual curation, thus providing practical benefits across biomedical research, clinical practice, pharmaceutical industries, and educational applications.

Transformer-based large language models (LLMs) have emerged as promising tools to address this need due to their impressive capabilities in information synthesis and natural language processing. Indeed, these models have demonstrated notable successes, such as passing medical licensing exams (USMLE)[17] and performing competitively in medical subspecialty assessments [18, 19, 20]. Despite these strengths, LLMs have notable limitations, particularly concerning inaccuracies in information retrieval and the generation of fabricated content (hallucinations). Studies by Bhattacharyya et al. (2023), Walters et al. (2023), and Chelli et al. (2024) highlight these issues, showing high rates of inaccuracies and fabricated references across various LLMs including GPT-3.5, GPT-4, and Bard [21, 22, 23].

Consistent with these concerns, our results identified similar limitations. ChatGPT-4o had 16.4 % hallucination rate, produced 18.4% topic-irrelevant citations, and achieved only 59.3% accuracy in gene-to-phenotype mapping. Gemini 1.5 Pro, another transformer-based LLM, demonstrated slightly lower mapping accuracy (55.8%) but produced fewer irrelevant citations and fabricated references, suggesting variability in performance among models.

The suboptimal accuracy of these general-purpose LLMs likely results from their inability to directly access the peer-reviewed articles, restricting their literature retrieval capabilities and accuracy in synthesizing gene-to-phenotype relationships [24]. Moreover, their inherent design for linguistic rather than information processing predisposes these models to generate inaccuracies and fabrications [22].

To overcome these limitations and improve practical applicability, specialized scientific LLMs have been developed. These specialized models incorporate enhanced features specifically designed to improve literature retrieval and information extraction accuracy, such as abstract retrieval (PubMedGPT), integration with academic databases (ScholarGPT), and augmentation with proprietary databases containing extensive collections of full-length articles (ScholarAI and SciSpace).

Performance varied notably among these specialized models. PubMedGPT, despite targeted access to PubMed abstracts and open-access full texts via PubMed Central, had moderate accuracy (57.6%) and low irrelevant citations (5.4%), but suffer from high hallucination rate similar to the base ChatGPT-4o model (16.2%). Conversely, models with extensive access to full-text articles (ScholarAI and SciSpace) markedly outperformed others, achieving accuracies of 81.1% and 74.9%, respectively, with the lowest rate of hallucination rate and irrelevant references among all the models tested in this study. These results underscore the critical role of comprehensive database access in enhancing LLM performance, with potential to benefit researchers through expedited literature reviews, clinicians via rapid candidate gene identification, pharmaceutical companies by streamlining drug target validation, and educational settings through authoritative learning resources.

Our findings extend and complement previous studies in this field. Toufiq et al. (2023) demonstrated LLM utility in knowledge-driven gene prioritization, albeit within predefined modules rather than direct gene-to-phenotype mapping [26]. Kim et al. (2024) found limited accuracy of GPT-4 in phenotype-driven gene prioritization for rare diseases, noting biases toward highly studied genes [27]. Our results expand upon these findings by demonstrating significantly improved accuracy when LLMs are augmented with comprehensive full-text databases, highlighting the importance of integrating extensive domain-specific knowledge sources for genomic applications.

Future research should build upon these findings through systematic optimization of prompt engineering, exploring strategies such as multi-shot prompting, chain-of-thought reasoning, and domain-specific prompt libraries to enhance accuracy and reliability. Additionally, exploring reverse phenotype-to-gene mapping, broadening the scope and scale of evaluation across diverse biological domains, incorporating temporal aspects of gene expression and phenotype development, and integrating with established databases like OMIM, ClinVar, and GWAS catalogs would significantly advance the applicability of these models. Furthermore, the development of standardized protocols for clinical integration, quality control, and interpretation frameworks is essential for practical, safe, and effective real-world use.

Despite these promising outcomes, our study has limitations, including the evaluation of a limited subset of existing specialized GPT models and relatively small sets of genes and phenotypes compared to the entire human genome. Future studies addressing these limitations through expanded evaluations will be vital for validating and generalizing our findings.

## Conclusion

This study represents the first systematic and comprehensive analysis of both base transformer-based large language models (LLMs) and specialized LLMs built on the GPT-4 architecture. Specifically, we evaluated their ability to retrieve query-relevant scientific literature and synthesize this information to generate accurate gene-to-phenotype mappings. The GPT-4-based LLMs, particularly when equipped with direct access to databases, demonstrated significant improvement in accurately mapping gene-to-phenotypes. This integration not only enabled the generation of highly accurate gene-to-phenotype mappings but also ensured that the outputs were supported by relevant, peer-reviewed scientific articles. By combining cutting-edge natural language processing capabilities with access to high-quality data sources, GPT-4-based LLMs highlight their potential as powerful tools for facilitating complex biomedical research tasks. This bridges the gap between vast genomic datasets and actionable biological insights

## Figures

**Supplementary Figure 1:**
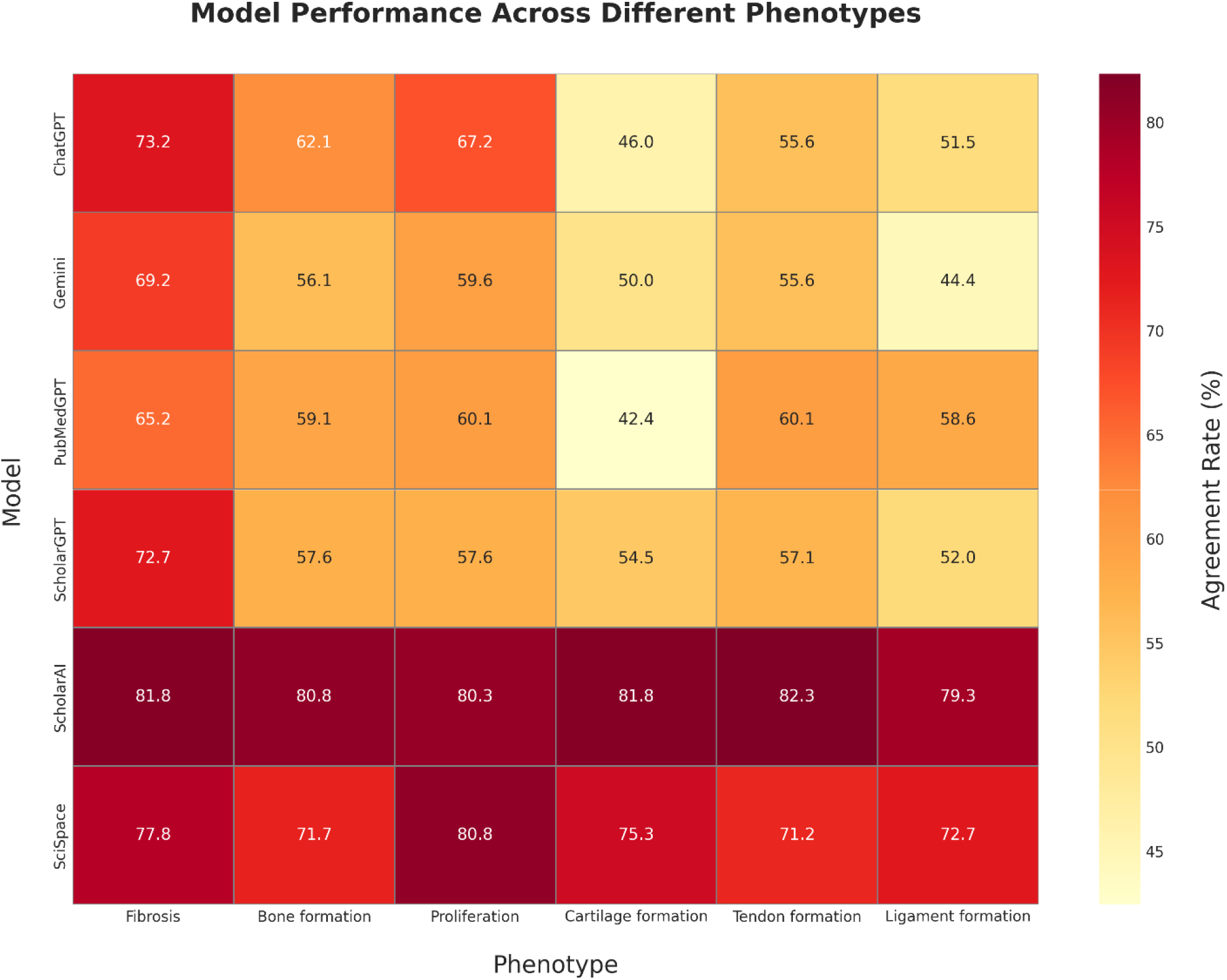
Phenotype-specific performance heatmap showing agreement rates (%) for each model across six different phenotypes. The heatmap reveals consistent patterns: cartilage formation (lighter yellow regions) poses the greatest challenge across all models, while fibrosis and cell proliferation (darker red regions) generally show higher agreement rates. Database-augmented models (ScholarAI and SciSpace) maintain more consistent performance across phenotypes compared to general-purpose LLMs, which show greater variability.

## Author Contributions

Nicolas Suhardi and Vincentius Suhardi selected the list of genes and phenotypes, tested the relevance and accuracy of the references provided by the LLMs, conducted data analysis, and formulated questions. Anastasia Oktarina, Damanpreet Dhillon, and Dona Ninan recorded the responses generated by all LLMs and organized the raw text responses into Excel spreadsheets. Anastasia Oktarina verified the relevance and accuracy of the references provided by LLMs. Vincentius Suhardi performed the manual curation of references and manual gene-to-phenotype mapping that are used as ground truth for this study. Mathias Bostrom and Xu Yang assisted in editing the manuscript.

## Data Availability

All data generated in this study are available from the corresponding author upon reasonable request.

## Acknowledgments

The authors thank the research staff at Hospital for Special Surgery for their support in this study. We acknowledge the contributions of the large language model developers who made their systems publicly accessible for research purposes.

